# Chiral Single Molecule Localization Microscopy (chiralSMLM)

**DOI:** 10.1101/2025.02.10.637296

**Authors:** S Aravinth, Neeraj Pant, Partha P. Mondal

## Abstract

Chiral fingerprints are unique to a molecule and convey information related to binding sites, conformation, and its immediate chemical environment. Deciphering this information is expected to aid a better understanding of the biological processes (protein trafficking, kinetics, and aggregation) in a cellular environment at the scale of a single molecule. Here, we present a new optical technique called *chiralSM LM*, to selectively detect chiral enantiomers (both left-handed and right-handed) of single molecules and generate a super-resolved map to understand the role of chirality associated with the biological processes in a cell. Accordingly, calibration, characterization, and optimization of the system are carried out using a polarized light source (linear polarized, left and right circularly polarized) and by recording known structures (Actin filaments) in a cell. The system is employed to carry out cell transfection studies on two different disease models (Hemagglutinin protein for Influenza type-A and NS3 protein for Dengue type-2) to understand the role of molecular chirality on disease specific biological processes leading to clustering. Single-molecule cluster analysis revealed that left-handed Dendra2-HA and Dendra2-NS3 molecules have a larger footprint, suggesting the role of chiral molecules in promoting cluster formation. In addition, the presence of left-handed molecules at the cluster-periphery is perplexing. This is interesting, as it demonstrates the active role of single molecule handedness (left or right) during protein clustering in a transfected cell. The new classification of single molecules purely based on their chiral nature is expected to advance single-molecule imaging and provide new insights in disease biology.

**Summary & Significance:** The chirality of single protein molecules is crucial to its functioning. This provides a new perspective on understanding biological functions, based solely on the chiral nature of proteins, and deciphers its role in a cellular processes. We developed a *chiralSM LM* microscopy system to access the role of chirality on the clustering behavior of viral proteins (HA for Influenza type-A and NS3 in Dengue type-2) in a cell. Hence, the technique offers a new approach to quantify chirality aided molecular interactions in disease biology.

## I. INTRODUCTION

For decades, molecular chirality has been known to play crucial roles in natural sciences [1] [2] [3]. In biology, the chirality of a molecule enables a better understanding of the biological processes associated with binding, conformation, and recognition of intermediates and catalytic products. This enhances our understanding of the functioning of stereoselective biomolecules in a cellular system. However, these molecules often occur in mixed form (racemic mixture), and hence, detection of the chirality of a single biomolecule is of great interest in a range of fields across all the sub-disciplines of biological sciences. The chirality at the single molecule level is hard to detect as compared to the bulk, calling for specialized optical detection methods. Therefore, a technique capable of deciphering chirality at a single molecule level is of great interest to cell biology, drug discovery, and pharmacology.

The availability of chiral information, in addition to its position and localization information, adds a new dimension to single-molecule imaging. In general, super-resolution map gives the location of single molecules as evident from prominent SMLM techniques (PALM, fPALM, STORM, DNA-PAINT), and its variants (BP-fPALM, IML-SPIM, ROSE, SMILE, POSSIBLE, *scanSM LM*, *corrSM LM*) [15] [16] [17] [19] [22] [20] [21] [23] [4] [5]. These techniques, along with other super-resolution techniques, hold the key to unraveling the nanoscopic world of single biomolecules [24] [25] [47] [48] [49] [50] [24] [25]. Although these techniques produce ultra-high resolution map of the distribution of single molecules in a cell, there are no known techniques that can determine their chirality based on fluorescence. Noting that the chirality of a molecule gives key information related to folding, binding, and immediate chemical information, this becomes essential to understand underlying biological mechanism [26] [27] [28]. For example, the three-dimensional structures of proteins depend on the homochirality of amino acids. One of the enantiomers may fit an enzyme (through a binding site or chiral groove), but the other may not [14]. Hence, the chirality of a biomolecule is significant in carrying out the basic functioning of proteins / amino acids. In fact, chirality exists in carbohydrates, DNA, organelles, tissues, and organs. Moreover, most of the amino acids are left-handed, and most sugars are right-handed [6] [7] [8]. The fact that the structure of a protein, its folding, and its biological functions are intrinsically linked directly exposes the role played by chirality. In addition, stability and conformational flexibility, which are linked to chirality, are necessary for cargo transport, enzymatic processing, regulatory functions, and communication [29] [30] [32] [33]. These processes have deeper repercussions in a number of diseases (Alzheimer’s disease (AD), Parkinson’s disease, and Type II diabetes) [34] [35] [36].

Circular polarisation luminescence (CPL) based spectroscopy and microscopy techniques are key for quantifying chiral molecules and its function in cellular systems [37] [38] [39]. The chiral molecules selectively absorb right- or left-circularly polarized light, and this is reflected in their emission pattern. Since CPL heavily depends on the absorbance of the sample, the techniques employing this property makes it insensitive for cell imaging. To date, chirality is explored in many fields ranging from cell biology to material science. For example, the technique is quite useful for the investigation of inorganic substrates generally used in material characterization [40] [41] [42]. Recently, CPL spectroscopy is found to be very useful for studying optically active Europium(III) which can be used as a contrast agent for cell imaging [37] [43] [44]. Moreover, wavelength-dependent left/right circular polarized emission from Europium(III) is observed and the same is used for chiral contrast imaging [45]. Despite these developments, circularly polarised emission microscopy studies are limited to bulk [46]. With the advancement of modern microscopy, it is necessary to develop microscopy techniques that can combine CPL with single molecule imaging.

In this article, we report the detection of chiral single biomolecules in a cell and use it to understand their clustering, which is a critical step toward virion maturation. This is enabled by a dedicated chiral detection system where fluorescence from a single molecule is separated based on its chirality. To separate chiral fluorescence, a combination of a quarter-wave plate and a polarizing beam-splitter is used. The whole detection sub-system is realized on a double-channel 4*f* configuration, and the detection of fluorescence (from the single-molecule enantiomers) is observed on a single detector chip. In addition, a series of filters and steering mirrors are used to filter and direct both the beam (left-circularly and right-circularly polarized) onto the same camera chip, thereby enabling simultaneous detection of both the enantiomers. Subsequently, the technique is calibrated, standardized, and validated using fluorescent beads, circularly polarized light source and known structures (Actin filaments) in a cell. Finally, the method is employed to understand the effect of chirality on the clustering of conjugated viral proteins (Dendra2-HA and Dendra2-NS3) in transfected cells.

## II. RESULTS

### A. Chiral SMLM System

To investigate the chiral characteristics (left- and right-handedness) of a single biomolecule and its role in clustering during viral pathogenesis, a new kind of super-resolution microscopy system (*chiralSM LM*) is developed. Fig. 1 shows the schematic diagram of the developed *chiralSM LM* system. The system has two major optical sub-systems: illumination and detection. The illumination consists of two lasers: an activation laser of wavelength 405 *nm* for activating the molecule and a second laser of wavelength 561 *nm* for excitation, causing fluorescence emission. The lasers are combined using a dichroic mirror (DMLP425R, Thorlabs) and directed to the electrically tunable lens (ETL) (Optotune, −1.5 to +3.5 diopters, EL-10-30-CI-VIS-LD-MV), which is placed in front of epifluorescence port of Olympus IX 81 inverted microscope. The ETL focuses the combined beam at the back aperture of the objective lens (Olympus, 100X, 1.3NA) using a dichroic mirror DM2 (Di03-R561-t1-25*36, Semrock) placed inside the filter cube of the microscope.

**FIG. 1:**
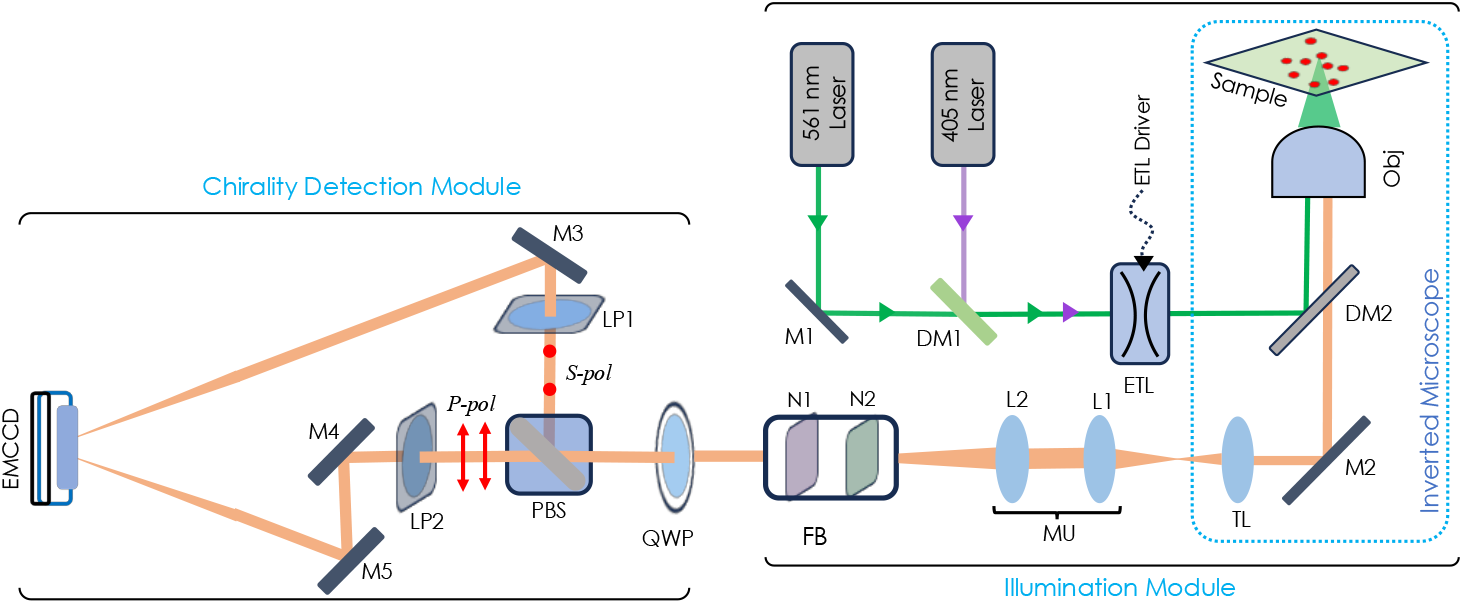
Schematic diagram of chiral SMLM: The figure illustrates the optical setup of chiralSMLM that consists of laser sources, inverted microscope and chiral detection module.

The chiral detection module consists of a Quater Wave Plate (QWP, 10RP54-1B, Newport) mounted on a precision cage rotation mount (CRM1PT, Thorlabs). The fast axis of the QWP is aligned 45° to the *s*-polarized light (*s*-polarization vector perpendicular to the optical table). This converts Right Circularly Polarized light (RCP) into *s*-polarized light and Left Circulary Polarized light (LCP) into *p*-polarized light. The *s*-polarized light is reflected by the polarizing beam splitter (PBS)(#49002, Edmund Optics), and *p*-polarized light is transmitted. This combination of QWP and PBS split the RCP and LCP fluorescence signal emitted by the single molecules into two paths. The mirror M3 directs the RCP light to the right side of the camera sensor (RCP channel), and the mirrors M4 and M5 direct the LCP to the left side of the camera sensor (LCP channel). The mirror *M*5 is placed on a linear translator stage, and it is moved precisely to ensure that the fluorescence signal on both channels is at a focal length of the tube lens (TL). The perfectly aligned linear polarizers (LP1 and LP2) ensure only *s* and *p* polarized light enters the camera. Also, the detection subsystem is equipped with two notch filters, N1, and N2 (NF03-405E-25,NF03-561E-25), placed in the filter box (FB) and a long pass filter LPF (561nm Long pass filter - LP02-561RE-25). These filters help eliminate the illumination lights (405 nm and 561 nm) and minimize the background, allowing fluorescence to reach the detector. Overall, the optical setup facilitates simultaneous recording of left and right circularly polarized fluorescence light emitted by left and right-handed single-molecules.

### B. Calibration and Standardization

The fluorescence signal from both the channels is collected simultaneously on a single camera chip placed at the focal length of the tube lens. Simultaneous detection requires precise calibration which is performed by imaging fluorescent nanobeads (175nm, Invitrogen *ex*: *em* | 505: 515*nm*) embedded in Agarose gel-matrix. First, the bead image is focused on the RCP channel (top portion of the sensor chip), which is followed by precisely changing the distance between M3 and M4 such that the focused image of the bead appears on the LCP channel (see, Fig. 2A). The center of the bead on each channel is calculated, and based on that, lateral shift along the X and Y direction of the RCP channel with respect to LCP is determined. However, for axial shift, the bead was slightly defocused using the microscope nob. This resulted in the defocused image of the beads on both channels, the image of which is captured by the camera (see, Fig. 2B1). This defocused image of a bead consists of airy disk pattern, and accurate matching of this pattern (along with the rings) ensures same optical path for both channels, since slight off-set will result in change in size and position of the rings. Subsequently, the intensity is plotted along a line that passes through the center of each airy disk, and the intensity curve is correlated. The correlation is found to be 94.61%, confirming that signal from both channels are at the same focal distance. The fast axis of linear polarizers is aligned using a linearly polarized laser source and a power meter. The LP1 mounted on the precision cage mount is kept on a laser source which emits s-polarized light, and the fast axis of LP1 is rotated in such a way that it transmits maximum laser light. However, for LP2 the fast axis is aligned for the same laser where the transmitted laser power is minimum, and this ensures the proper alignment of linear polarizers to pass *s*- and *p*-polarized light. To ensure that the top portion of the camera sensor captures RCP light and the bottom portion captures LCP light, the quarter-wave plate (QWP) is calibrated using circularly polarized light generated by placing a circular polarizer in front of the linearly-polarized laser light source (see, Fig. 2A). The fast axis of QWP is rotated to get the maximum intensity in the top window and the image is recorded by the camera (see, Fig. 2.B2). Later, the right circular polarizer is replaced with the left circular polarizer and this time, the bottom window shows higher intensity compared with the top window. This ensures that the top window records the RCP light whereas the bottom window records LCP light, and this is also confirmed by the contrast between two windows in both cases, i.e, 90.76 ± 0.4% and 91.78 ± 0.6%, respectively. Also, the contrast between two windows is 2.11 ± 0.26% when only linear polarizer is used.

**FIG. 2:**
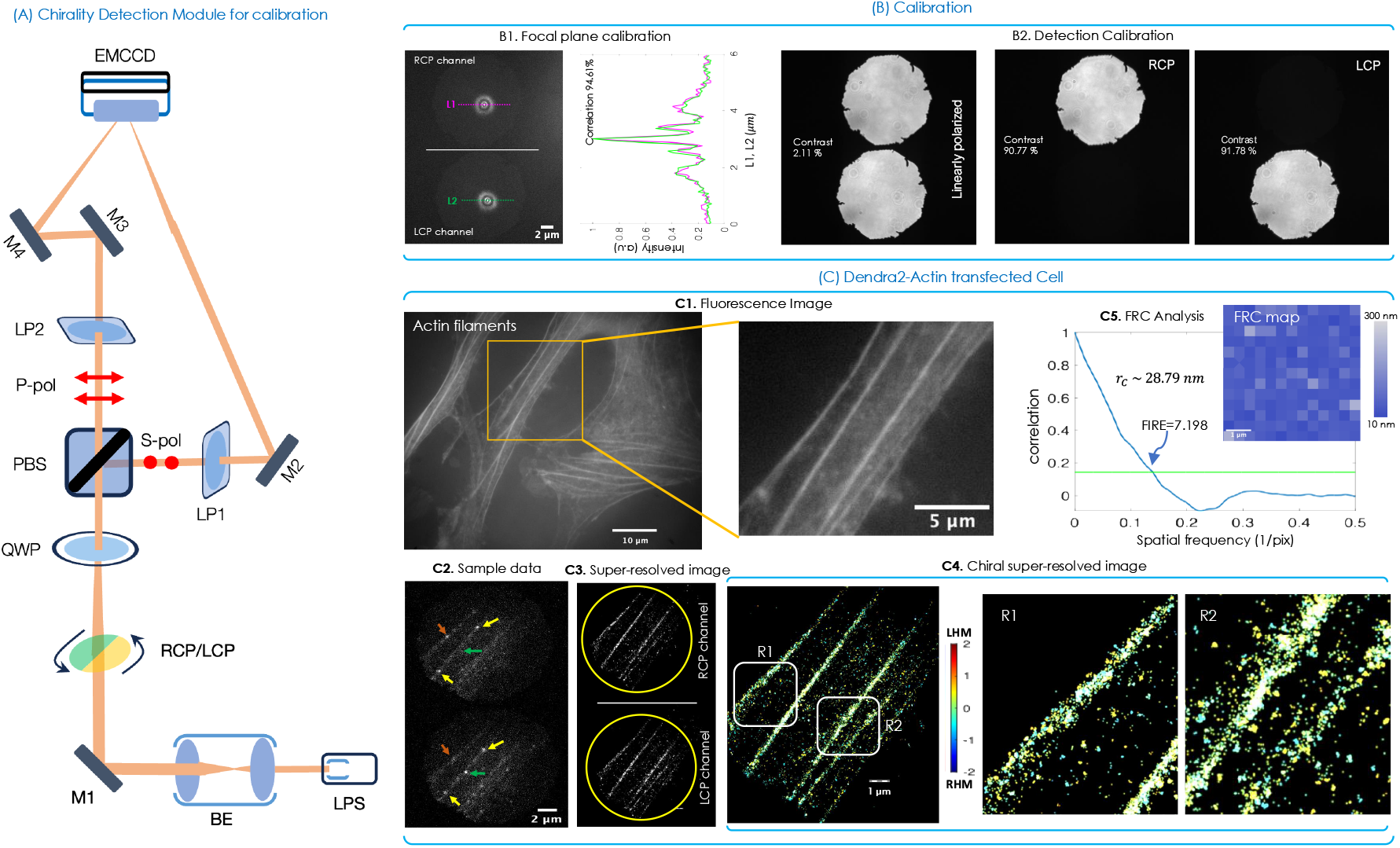
Chiral detection module, system calibration, quantification and image resolution: (A) Graphical representation of detection module of *chiralSM LM* for calibration. (B1) The focal plane calibration of *chiralSM LM* system is performed by imaging the fluorescence beads on both the channels. The figure shows the PSF of fluorescent beads in both the channels, and intensity is plotted for the PSF along a line passing through the center (magenta and green line) in each channel. The higher correlation(94.61%) of intensity plots confirms that the signal is recorded from a same focal plane. (B2) The detection module is calibrated using circularly polarized light. The top channel is brighter when right-circular polarizer is used, and the bottom channel is brighter when left-circular polarizer is used. Both the channels have equal intensity for linearly-polarizd light. The contrast between the top and bottom channels are mentioned. (C1) Fluorescence image of a transfected cell is recorded using fluorescence microscope and super-resolution study is carried out on a marked region(yellow box). The fluorescence image of the Dendra2-Actin transfected NIH3T3 cell and enlarged view of the cell is shown where, super-resolution imaging is performed. (C2) Raw data of single molecule blinks shows that single molecule PSF intensity is different with respect to the molecule in two channels as shown by yellow, green and orange arrows. (C3) The reconstructed super-resolution image of RCP and LCP channels. The chiral dissymmetry factor (*g*) is calculated, and the molecules are color-coded based on their dissymmetry factor value and presented as chiral super-resolved image (C4). Two enlarged regions(R1 and R2) of chiral super-resolution image are shown. (C5) The Fourier Ring Correlation analysis. This gives resolution of the chiral super-resolved image (~ 29 nm). Also, the FRC map for the chiral super-resolution image is generated using SQUIRREL analysis, and it is shown with the calibration bar. The corresponding raw data for Dendra2-Actin is shown in **Supplementary Video**.

### C. Actin Filaments in NIH3T3 Cells

The technique (*chiralSM LM*) is tested on known biological samples with well-known structures such as Actin filaments. Fig. 2(C) shows the result obtained for Dendra2-Actin transfected NIH3T3 cells. The transfection protocol is discussed in the Methods section. The filaments are imaged in a transfected cell to determine the fraction of chiral Actin molecules / proteins. Initially, the transfected cell image is captured using high-resolution fluorescence microscope, as shown in Fig. 2(C1), and a small region of the transfected cell (marked by yellow box) is imaged using *χ*-*SM LM*. A sample frame of the recorded data is shown in Fig. 2(C2), which clearly indicates the intensity-difference between the channel for a sample Dendra2-Actin molecule as marked by arrows. The corresponding reconstructed super-resolved images of the left- and right-handed Dendra2-Actin molecules (represented by chiral dissymetry factor, *g*) along with the merged image is shown in Fig. 2(C3,C4). The chiral dissymmetry factor (g) of each molecule is calculated using, 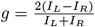, where *I*_*R*_ and *I*_*L*_ are the intensity of the molecules in the RCP and LCP channels respectively. Fig. 2(C5) shows the FRC analysis along with the FRC map [54] [55]. The reconstructed images have a Fourier Image Resolution(FIRE) of 7.198, which corresponds to a resolution (*r*_*cs*_) of 28.79 *nm*. The analysis clearly indicates high resolution along with the chiral footprint (see, Fig. 2C3, colormap image) of single molecules.

### D. Cell Transfection Studies: Disease Biology

Two different disease models are chosen and the chiral distribution of single viral proteins are investigated. NIH3T3 cells were transfected with photoactivable viral proteins (Dendra2-HA, cloned viral protein for Influenza type-A and Dendra2-NS3, a cloned viral protein for Dengue type-2) using the developed protocol (see, methods section). The cells were fixed for imaging after 48 hrs of transfection. The single molecule data is collected using the developed *chiralSM LM* microscope, and the images are reconstructed along with chiral information color-coded on each molecule (green for LCP with *g* ∈ [+0.2, +2] and magenta for RCP with *g* ∈ [−0.2, −2]) as shown in Fig. 3A. The combined image along with the enlarged section of few clustered regions are also displayed. The super-resolved chiral images shows the presence of left-handed molecules (magenta) at the periphery of few clusters, suggesting the role of single molecule chirality during cluster formation. To further substantiate, we carried out chirality analysis of HA and NS3 clusters. For this, we considered the number of molecules present in a cluster and calculated chiral ratio (*ζ* = *N*_*g*<−0.05_*/N*_*g*>0.05_) of each cluster. The molecules in the window *g* ∈ [−0.2, +0.2] is left out as they are the ones which exhibit low chirality. Note that, a cutoff of | 0.2| is chosen for the study. Few clusters, *R*1 − *R*4 and *S*1 − *S*4 in Dendra2-HA and Dendra2-NS3 transfected cells are displayed along with the *ζ*-value. Alongside, the chiral ratio plot of all the clustered molecules is shown in Fig. 3B, indicating the average chirality from the total number of molecules participating in cluster formation. However, a separate chiral histogram plot is added that shows clusters-wise chirality, indicating the dominance of left-handed (LH) molecules.

**FIG. 3:**
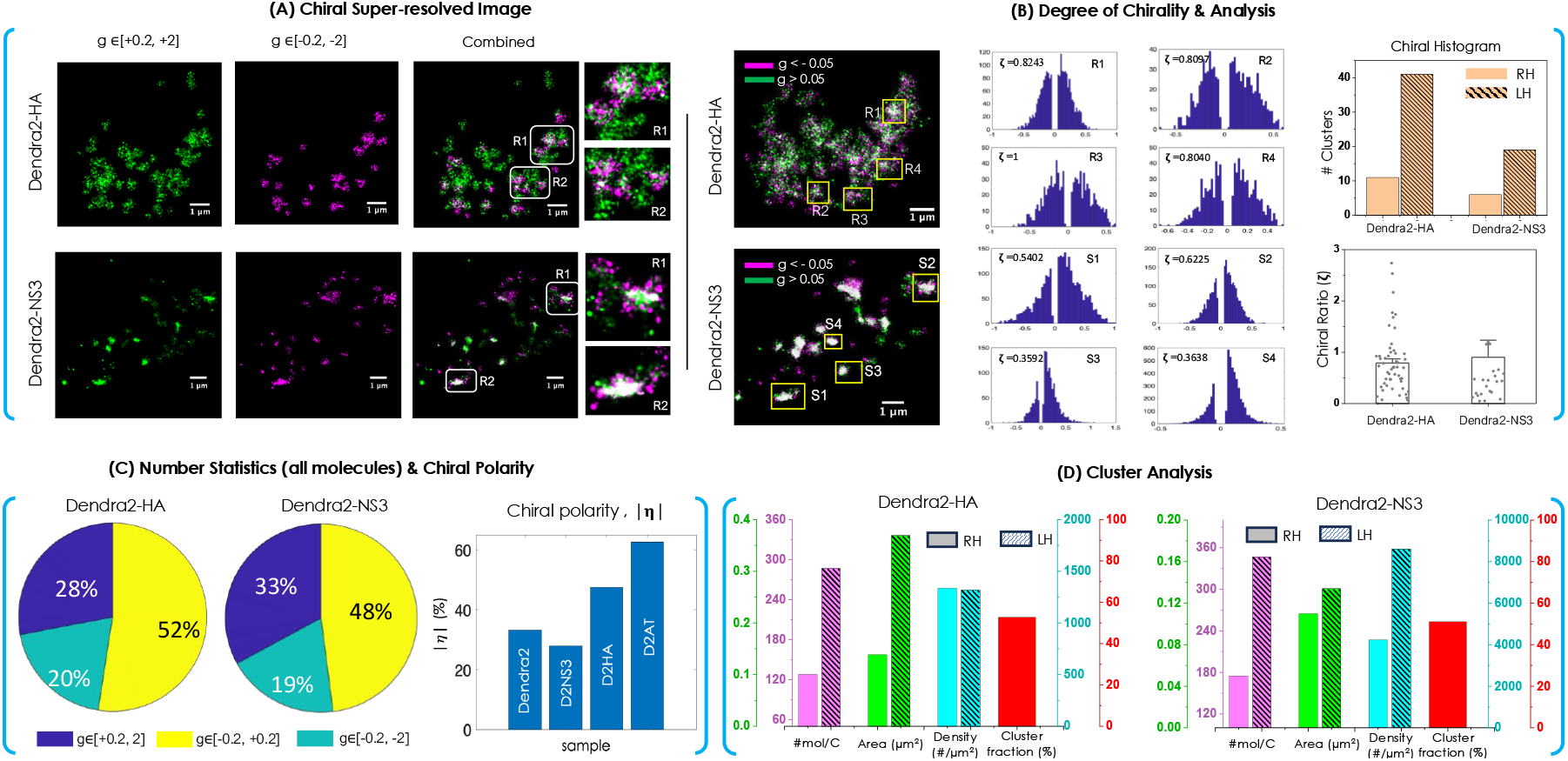
Dendra2HA and NS3 images: (A) The chiral super-resolved images (Dendra2-HA and Dendra2-NS3 transfected cells) are shown. Color code is used to represent the chirality of single molecules, with green for *g* ∈ [+0.2, +2], and magenta for *g* ∈ [−0.2, −2]. Enlarged sections of the combined images show the distribution of chiral molecules in the single molecule clusters. (B) Chirality based analysis of the clusters (HA and NS3). Single molecule histogram for a few chosen clusters (R1-R4, and S1-S4) are displayed. Alongside, the chiral histogram of the clusters (indicating overall chirality of individual cluster) and chiral ratio (*ζ* = *N*_*g*<−0.05_*/N*_*g*>0.05_) plot for the number of molecules forming clusters is also shown. This indicates a predominant left-handed nature of HA and NS3 clusters. (C) The pie chart shows the percentage of molecules with, *g* ∈ [−0.02, −2], *g* ∈ [−0.2, +0.2], and *g* ∈ [+0.2, +2] considering all the detected molecules. In addition, chiral polarity (|*η*| = *Ng* > |0.2| */N*_*T*_) for all the molecules is also displayed. (D) The average cluster-related parameters calculated using DBSCAN for both Dendra2-HA and Dendra2-NS3 are shown in the bar graph along with cluster fraction (the percentage of molecules forming clusters). The corresponding raw data for Dendra2-HA and Dendra2-NS3 can be found in **Supplementary Video**.

Next, we focus our attention on the overall number statistics of all the molecules (both participating in cluster formation and otherwise). The pie-chart shown in Fig. 2C for Dendra2-HA clearly suggests a good fraction (28%) of left-handed (*g* ∈ [+0.2, +2]) as compared to right-handed (*g* ∈ [−0.2, −2]) molecules (20%). However, a large fraction 52% still does not participate in cluster formation. This same signature is observed for Dendra2-NS3 transfected cells for which the left-handed is about 33% of the total detected molecules. A histogram is plotted to determine the total number of chiral molecules (molecules with chirality |*η*| > 0.2). The chiral polarity (*η*) is discussed in the methods section. It is evident from the histogram that Dendra2-Actin (D2AT) has more chiral molecules as compared to D2HA and D2NS3. However, the chiral molecules are randomly distributed for D2AT, whereas they are largely concentrated in the clusters for D2HA and D2NS3. Finally, cluster analysis is carried out for the molecules participating in clustering and exhibiting chiral nature (RH or LH). The analysis comprises of the average number of molecules in a cluster, the average area of cluster, and its density. Alongside, the cluster fraction (percentage of chiral molecules participating in clustering) is also shown. On an average, LH molecules are present in large numbers in clusters, and have larger area / spread. However, they (LH and RH molecules) have comparable density for Dendra2-HA and relatively large density for Dendra2-NS3. The analysis overall indicates larger footprint of left-handed molecules as compared to right-handed molecules and chirality has a significant role to play during cluster-formation stage post infection (here, post 48 hrs of transfection). The study also highlights the role of chirality in promoting clustering in cell transfection studies (Influenza type-A and Dengue type-2). The observation may have implications in other viral disease falling in the broad family, flavi-virade.

## III. DISCUSION & CONCLUSION

Chirality is an intrinsic property of a protein molecule and an important cofactor that regulates protein functioning in a cell. This is central to maintaining cell physiology during viral infection. Here, we present a new technique to optically detect single-molecule chirality, where the emitted fluorescence from single molecules is split into *RCP* and *LCP* components to separate out enantiomers. The method is compatible with the standard SMLM systems and blends well with the existing image analysis algorithms. However, dedicated dual-channel 4*f* optical system equipped with specialized optical elements in the detection module is a must. To calibrate a known polarized light source (linear, LCP and RCP light) is employed. In addition, to test the performance of *chiralSM LM*, known cell structure (Actin filaments in NIH3T3 cells) is used. The molecules are studied, based on their chirality: RH (*g* ∈ [−0.2, −2]), and LH (*g* ∈ [+0.2, +2]). Subsequently, a super-resolved image is reconstructed by combining molecules with negative (left-handed) and positive (right-handed) chirality.

The *chiralSM LM* system is used to understand the role of chirality in disease models: Influenza type-A, and Dengue Type-2. Transfection cell studies reveal the formation of single-molecule / protein clusters. The results show a relatively dominant presence of left-handed molecules in clusters with a large footprint during viral infection, especially these molecules appeared at the periphery of a few Dengue NS3 clusters. We could not determine the peculiar behavior behind the presence of left-handed-NS3 molecules at the periphery of these clusters. Analysis shows a significant fraction of the molecules have a chiral nature (61% for Dendra2-Actin, 45% for Dendra2-HA, and 27% for Dendra2-NS3). Single-molecule cluster analysis exemplifies this, with left-handed molecules dominating cluster formation, and exhibiting specific characteristics with respect to the number of molecules per cluster, cluster spread/area, and cluster density. This is unique for each case, i.e, Dendra2-HA and Dendra2-NS3.

Overall, the newly developed *chiralSM LM* system exemplifies the chiral nature of single molecules and their significance in disease biology. The clusters thus formed have a dominant presence of left-handed molecules, which may have a role in promoting and stabilizing clusters. However, the mechanism behind such behavior of molecular handedness is not understood well and may require additional studies to assess the consequences. Nonetheless, the new optical microscopy technique and the related discovery (chirality-promoted clustering) can have therapeutic significance, with potential drugs disrupting the molecular clusters.

## IV. METHODS

### A. Cell culture and Transfection

NIH3T3 are seeded on No.1(22mm x 22mm) coverslip with a cell count of 90,000. 16 hours post seeding, once the cells are attached properly, the transfection is performed using Lipofectamine 3000 kit(ThermoFisher Scientific, Catalog number:L3000008). The DNA complex is prepared in Opti-MEM media(ThermoFisher Scientific, Catalog number: 11058021). 1*µg* of plasmid DNA 2.25*µL* of p3000 and 3*µL* of Lipofectamine was used. 48 hours post transfection cells were washed with 1x PBS two times and fixed with 3.7% of paraformaldehyde. Then, the coverslip is mounted on the glass slide with Fluorosave mount media.

### B. Bead sample preparation

20*mg* of Agarose is mixed with 5*mL* of milli-Q water and heated using a hotplate at ~ 100°*C*. Once Agarose dissolved in water, the hotplate is turned off, and the temperature of the solution is allowed to decrease. When the temperature is close to 40°C 1*µL* of bead solution is added to the Agarose solution and uniformly mixed. Then, the mixed solution is casetted on the live imaging dish.

### C. Data Collection and Analysis

EMCCD camera is used to record the fluorescence signal from the single molecules. Before data collection, the temperature of the camera is cooled to −88°*C* to avoid thermal noise. The frame transfer mode is enabled, and EM gain is kept at 270. The data is recorded in ‘.tif’ file format with an exposure time of 40*ms*. The recorded data were analyzed using lab-made MATLAB program, which involves background subtraction using a rolling ball algorithm followed by single-molecule spot detection using the intensity threshold method. The detected spots are fitted with the 2D Gaussian function based on the non-linear least square method. The standard deviation and centroid are measured from the fitted Gaussian, and also the number of photons is calculated. The localization precision of each molecule is calculated using the Thompson formula[52].

For chiral molecule analysis, the molecules detected in the bottom window is separated, and their coordinate is shifted based on the beads data then compared with the molecules detected in the top window of the same frame. Initially, the distance between the centroid of the molecules in both the windows after shift correction is calculated, when the centroid of the molecules is within 120*nm*, they are considered as same molecule and chiral dissymmetry factor is calculated based on their number of photons. The super-resolution image is reconstructed by collating all the molecules into a single frame, and each molecule is represented as 2D Gaussian function with the standard deviation of 20nm. For chiral super-resolution image the dissymmetry factor information is incorporated in the molecule color.

### D. Chiral Polarity

Chiral polarity is defined by,

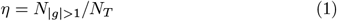

where, *N* _|*g*| >1_ is the number of molecules with |*g*| > 1, and *N*_*T*_ is the total number of molecules.

### E. Cluster Analysis

To identify clusters, DBSCAN clustering algorithm is used. DB-SCAN is a density-based technique that identifies groups points that are closely packed together in data space. The inbuilt Matlab function ‘*dbscan*’, along with machine learning toolbox is used to analyze singlemolecule clusters. The X and Y coordinates of the single detected single molecules are given as input, and cluster parameters such as the minimum cluster point (~ 50 points) and the minimum distance (2 pixels = 120nm) are chosen for the study. Once the molecules are clustered, the number of molecules per cluster, area of the cluster, and cluster density (number of molecules per unit area) are calculated.

### F. Optical Setup

The illumination subsystem of *chiralSM LM* is similar to *aSM LM* [53] where the ETL is operated at 270 *mA* current to get a required field of view. The specimen is illuminated with an intensity of 24.4*mW/µm*^2^ for activation and 93.3*W/µm*^2^ for excitation. In the detection subsystem, additional magnification is obtained using magnification unit (MU) which consists of two convex lens of focal length 150*mm* and 400*mm*, giving a magnification of 2.66*X*. Along with the 100*X* objective lens, the total magnification is about 266*X*. This ensures the oversampling of single molecule PSF. Given the pixel-size of 16 *µm* of the EMCCD camera, and a magnification of 266, the effective pixel size is 60*nm*.

## Acknowledgments

The authors acknowledge parent institute (Indian Institute of Science, Bangalore, India) for financial support. The authors thank Dr. Jiby Mary Varghese for the Dendra2-NS3 recombinant plasmid preparation.

## Author Contributions

PPM and AS conceived the idea. AS, NP and PPM carried out the experiments. AS and NP prepared the samples. PPM wrote the paper by taking inputs from all the authors.

## Data Availability

The data that support the findings of this study are available from the corresponding author upon request.

